## Supplemental Figures for "Recording γ-secretase activity in living mouse brains"

[#] Equally contributed
* Corresponding authors

**Figure 1—figure supplement 1**

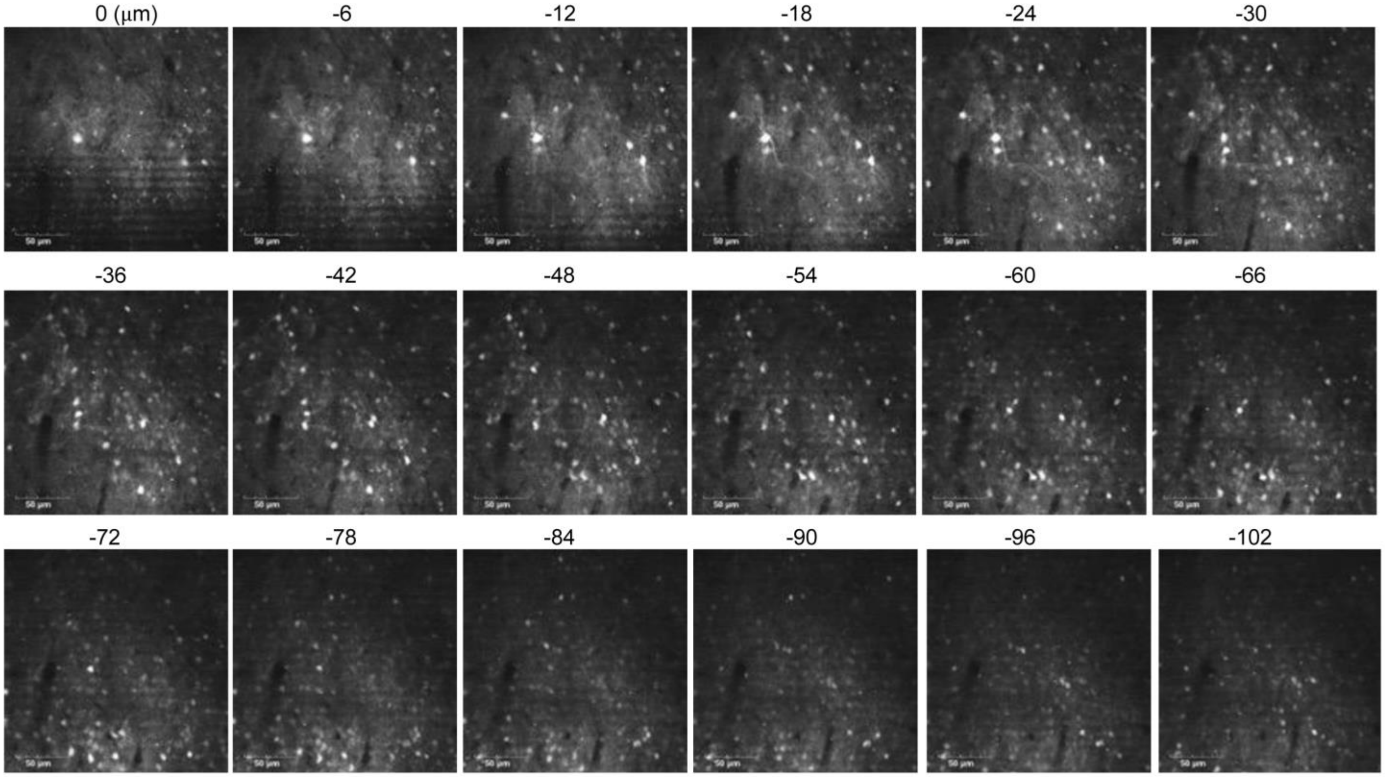


Z-section images of the brain expressing the C99 720-670 biosensor. The C99 720-670 biosensor fluorescence signal was detected at approximately 100 μm depth from the brain surface. Scale bar: 50 μm.

**Figure 1—figure supplement 2**

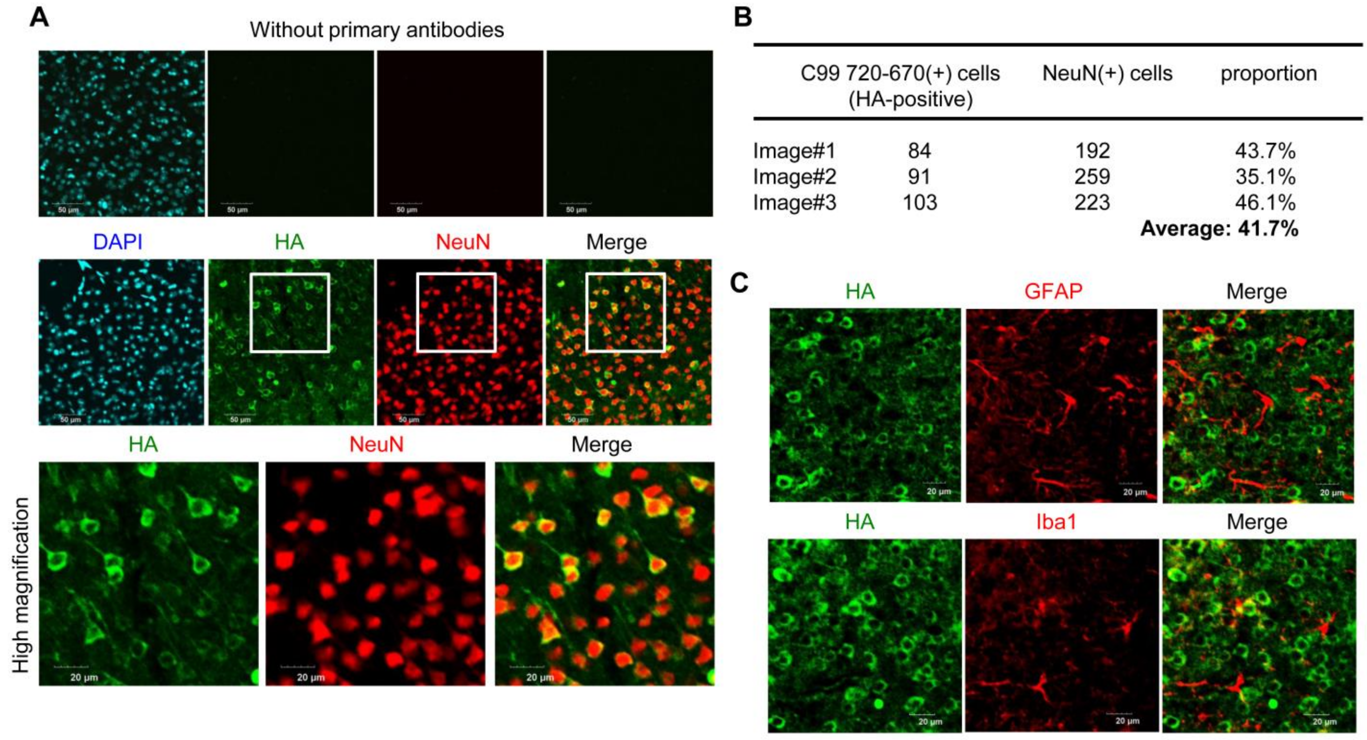


| | C99 720-670(+) cells (HA-positive) | NeuN(+) cells | proportion |
|---|---|---|---|
| Image#1 | 84 | 192 | 43.7% |
| Image#2 | 91 | 259 | 35.1% |
| Image#3 | 103 | 223 | 46.1% |
| | | | **Average: 41.7%** |

<u>Immunohistochemistry of the brain expressing the C99 720-670 biosensor</u>. **(A)** The brain sections of mice injected with an AAV-C99 720-670 were co-stained with HA and NeuN antibodies, showing that nearly 100 % of the cells expressing the C99 720-670 biosensor are NeuN-positive. Scale bar: 20 μm. **(B)** The quantification from three independent images suggests that approximately 40% of NeuN-positive neurons express the C99 720-670 biosensor. **(C)** Furthermore, we ensured that GFAP or Iba-1-positive cells do not co-localize with the HA signal, suggesting that the C99 720-670 biosensor is predominantly expressed in neurons. Scale bar: 20 μm.

**Figure 2—figure supplement 1**

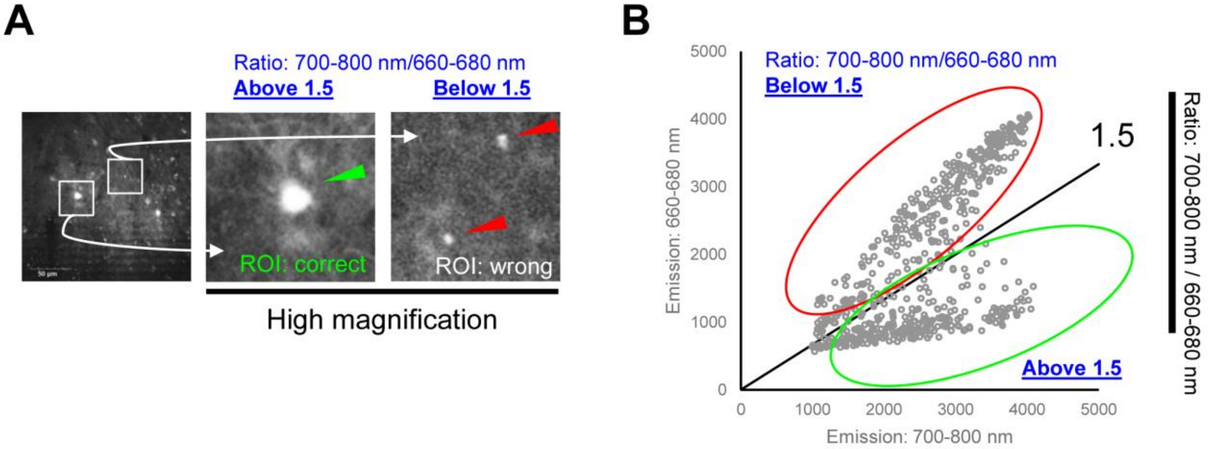


<u>Identification and removal of auto fluorescent objects</u> **(A)** Our four-step segmentation approach still could not perfectly remove wrongly assigned ROIs, such as shown in the right panel (red arrowheads). However, these ROIs displayed higher 660-680 nm emissions and thus significantly lower 720/670 ratios (below 1.5). **(B)** A scatter plot supported our observation**:** two populations of ROIs displaying 720/670 ratios above 1.5 and the ratios below, later of which were excluded from our data analysis.

**Figure 2—figure supplement 2**

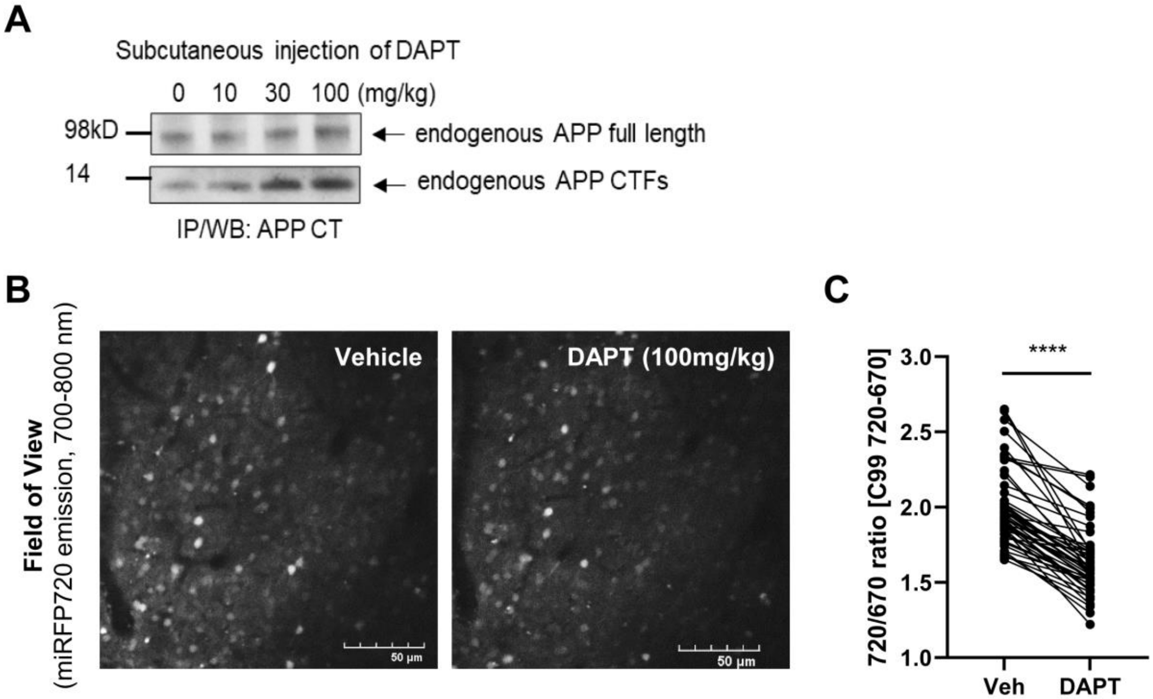


Validation of the C99 720-670 biosensor in the brain using γ-secretase inhibitor **(A)** The dose-dependent accumulation of endogenous APP C-terminus fragments (APP-CTFs) by subcutaneous administration of DAPT evidences the inhibition of γ-secretase activity in mouse brains. **(B** and **C)** The 720/670 ratios in the same neurons were compared before (vehicle) and 12 hours post 100 mg/kg DAPT administration. The 720/670 ratios were significantly decreased by DAPT administration (22.1%). N = 50 neurons, Mann-Whitney U-test, **** $p < 0.0001$.

**Figure 3—figure supplement 1**

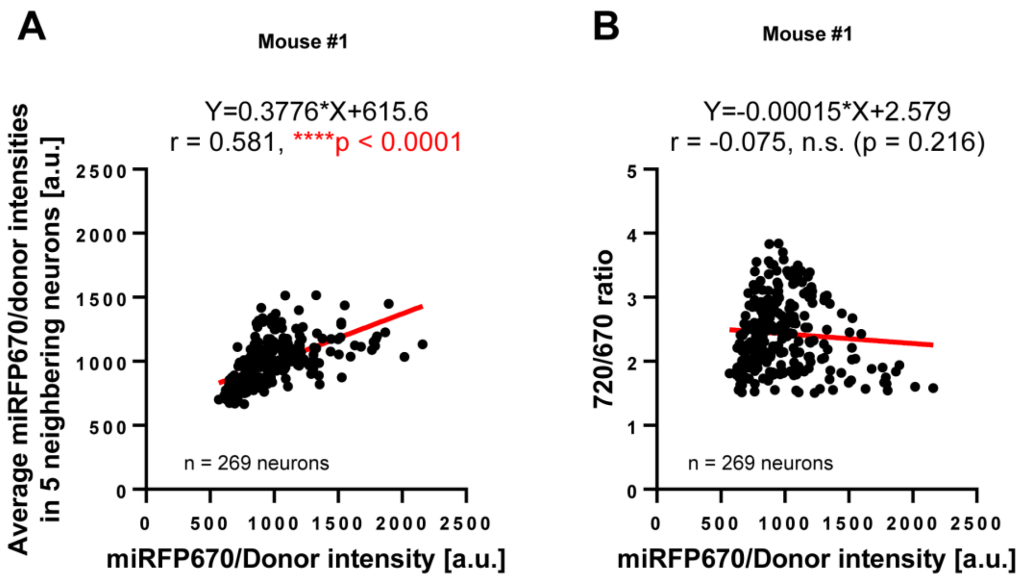


Expression pattern of the C99 720-670 biosensor **(A)** Scatter plots showing miRFP670 emission (as an indicator of the C99 720-670 biosensor expression) in individual neurons (X-axis) and the average Mean of miRFP670 emission in five neighboring neurons (Y-axis), suggesting uneven transduction of the AAV. The number of neurons, correlation coefficient (r), and p-value are shown. Pearson correlation coefficient. **** $p < 0.0001$ **(B)** However, the 720/670 ratio (i.e., $\gamma$-secretase activity) is not correlated with miRFP670 fluorescence intensity (i.e., C99 720-670 biosensor expression) in individual neurons. $p = 0.216$

**Figure 3—figure supplement 2**

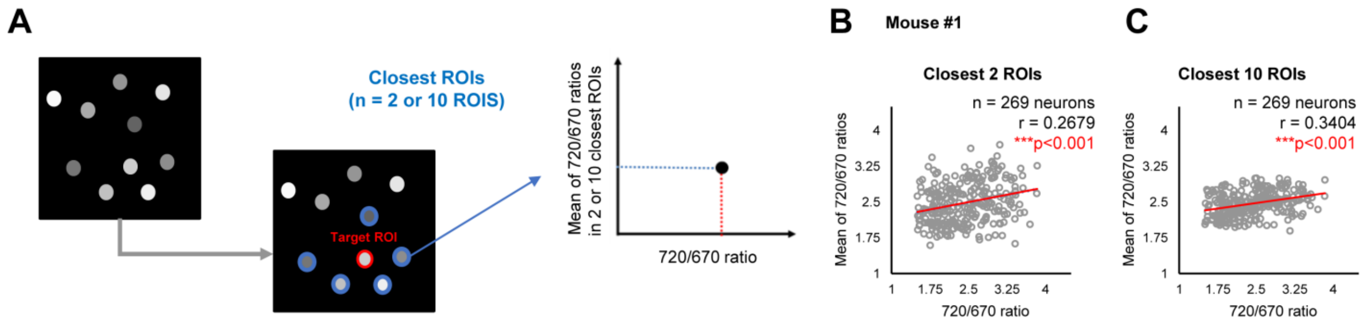


<u>Validation #1 - A potential "cell non-autonomous" regulation of γ-secretase in mouse brains</u> **(A)** The two or ten closest neurons were identified, and the average 720/670 ratio of the two or ten neighboring neurons was calculated and plotted. **(B)** We verified a significant positive correlation between the 720/670 ratio and the average ratio of two and **(C)** ten closest neurons. The number of neurons, correlation coefficient (r), and p-value are shown. Pearson correlation coefficient. *** $p < 0.001$
